# Dopamine D1-receptor Organization Contributes to Functional Brain Architecture

**DOI:** 10.1101/2023.03.24.534086

**Authors:** Robin Pedersen, Jarkko Johansson, Kristin Nordin, Anna Rieckmann, Anders Wåhlin, Lars Nyberg, Lars Bäckman, Alireza Salami

**Affiliations:** Department of Integrative Medical Biology, Umeå University, S-90197 Umeå, Sweden; Wallenberg Center for Molecular Medicine (WCMM), Umeå University, S-90197, Umeå, Sweden; Umeå Center for Functional Brain Imaging (UFBI), Umeå University, Sweden; Aging Research Center, Karolinska Institutet & Stockholm University, Tomtebodavägen 18A, S-17165, Stockholm, Sweden; Department of Radiation Sciences, Umeå University, S-90197 Umeå, Sweden; Max-Planck-Institut für Sozialrecht und Sozialpolitik, Amalienstrasse 33, 80799 Munich, Germany

## Abstract

Decades of research on functional brain mapping have highlighted the importance of understanding the functional organization of the cerebral cortex. Recent advances have revealed a gradient of functional organization spanning from primary sensory to transmodal cortices. This gradient-like axis of connectivity has been hypothesized to be aligned with regional differences in the density of neuromodulatory receptors. Recent work in non-human primates supports this notion, revealing a gradient of dopamine D1-like receptor (D1DR) density along the cortical hierarchy. Given the importance of dopaminergic modulation for synaptic activity and neural gain, we tested whether D1DRs shares the same organizational principles as brain function in humans, and whether inter-regional relationships in D1 expression modulate functional crosstalk. Using the world’s largest combined dopamine D1DR-PET and MRI database, we provided empirical support for the first time in humans that the landscape of D1DR availability follows a unimodal-transmodal cortical hierarchy, with greater D1DR expression in associative cortical regions. We found an organization of inter-regional D1DR co-expression spanning unimodal to transmodal brain regions, expressing a high spatial correspondence to the principal macroscale gradient of functional connectivity. Critically, we found that individual differences in D1DR density between unimodal and transmodal regions was associated with greater differentiation of default-mode and somatosensory networks. Finally, inter-regional D1DR co-expression was found to modulate couplings within, but not between, functional networks. Together, our results show that D1DR co-expression provides a biomolecular layer to the functional organization of the brain.

**Significance Statement:** We found a high correspondence between the organization of the most abundantly expressed dopamine receptor subtype and a macroscale unimodal-to-transmodal functional gradient. Differences in D1 density between unimodal and transmodal regions were related to the shape of the functional gradient, contributing to greater differentiation of somatomotor and default mode networks. Finally, we observed that the covariance structure of dopamine D1 receptors is associated with the strength of connectivity within functional networks. The discovery of a dopaminergic layer of brain organization represents a crucial first step towards an understanding of how dopamine, with close ties to behavior and neuropsychiatric conditions, potentially contribute to the emergence of functional brain organization.

## Introduction

The functional organization of the brain is assumed to be intrinsically related to cerebral microstructure. However, mapping between coordinated brain activity across distributed brain regions and their structural underpinnings have revealed a non-uniform structure-function tethering across the cortex ^1^, characterized by a gradual dissociation from unimodal to transmodal association cortices ^2,3^. A potential mechanism mediating the dissociation between structure and function is the organization of neurotransmitter systems. In particular, neuromodulatory transmitters may change the biophysical properties of action potentials ^4,5^ to integrate neural signals across spatially segregated brain structures ^6,7^. Indeed, recent work have shown that the spatial similarity between different neuroreceptor systems covary with structural pathways and moderate couplings between structural and functional connectivity ^8^. However, how the organization of individual receptor profiles may support functional architecture and modulate functional interactions is still poorly understood.

The neuromodulator dopamine (DA) plays an important role for synaptic and neural activity ^9,10^, and is associated with multiple physiological functions, including motor control, reward mechanisms, reinforcement learning, and higher-order cognition^11–14^. Human in-vivo imaging studies have revealed that DA D1 and D2-like receptors (D1DR and D2DR) are organized by distinct subsystems, reflecting anatomical differences between dopaminergic midbrain projections^15,16^ and functional systems^16,17^. Moreover, recent work in non-human primates have revealed a gradient in D1DR density along the cortical hierarchy^18^, characterized by greater receptor density in the associative cortex compared to somatosensory cortices. This pattern mimics the principal organization of cortical function, characterized by gradual differentiation in connectivity patterns from unimodal to transmodal regions^19,20^. Importantly, individual differences in D1DR and D2DR availability have been found to influence the strength of functional couplings^21–24^. It is therefore likely that the spatial arrangement of DA receptors contributes to large-scale functional architecture. However, it is not known whether the D1DR system, the most abundant DA receptor, expresses similar organizational properties to the functional connectome, and whether the spatial composition of D1 receptors modulates the topology of large-scale functional systems.

Using the world’s largest combined D1DR-PET and MRI dataset to date from the DyNAMiC study^25^, we set out to test the hypothesis that regional differences in D1DR density is related to the shape of the functional connectome and modulate the strength of functional couplings. To test the correspondence between functional and molecular organization, we first employed a non-linear embedding approach^26^ on group-representative covariance maps to effectively decompose complex spatial interactions into a more parsimonious set of organizing principles. In this framework, functional and dopaminergic organizations are characterized as a set of low dimensional manifolds, describing transitions in covariance patterns along the cortical surface^19,27–29^. Next, we extended our analyses to individual participants to assay whether the spatial differences in D1DR density that constitutes the molecular manifold accounts for inter-individual variation in functional organization, as indicated by differences in the relative position of regions in the functional manifold. Given the role of DA for functional distinctiveness^30^, we hypothesized that individuals with greater hierarchal differentiation in D1DR density express greater bimodality of the functional gradient, with a greater range between the gradient anchors. To not restrict our investigation to the low dimensional representations, we further investigated molecular-functional correspondence of regional interactions, capitalizing on discrete network boundaries. To this end, we used covariance matrices of each modality to investigate inter-regional associations between spatial D1DR covariance and functional couplings within and between canonical resting state networks.

## Results

We began by constructing a cortical profile of functional connectivity and D1DR organization based on rsfMRI and [11C]SCH-39166 PET images from a total of 180 healthy participants (for details on cohort characteristics, processing, and quality control, see SI Materials and methods) using an Laplacian eigenmapping^26^ approach. In brief, Laplacian eigenmapping is a nonlinear manifold algorithm able to resolve low-dimensional representations of spatial connectivity patterns commonly referred to as gradients. To this end, normalized functional connectivity matrices of 400 contiguous cortical parcels^31^ were decomposed into Laplacian eigenvectors. The values of each eigenvector represent the relative position of parcels in the embedding space, and distances between parcel positions indicate similarity in covariance patterns (Fig. 1).

**Fig 1.**
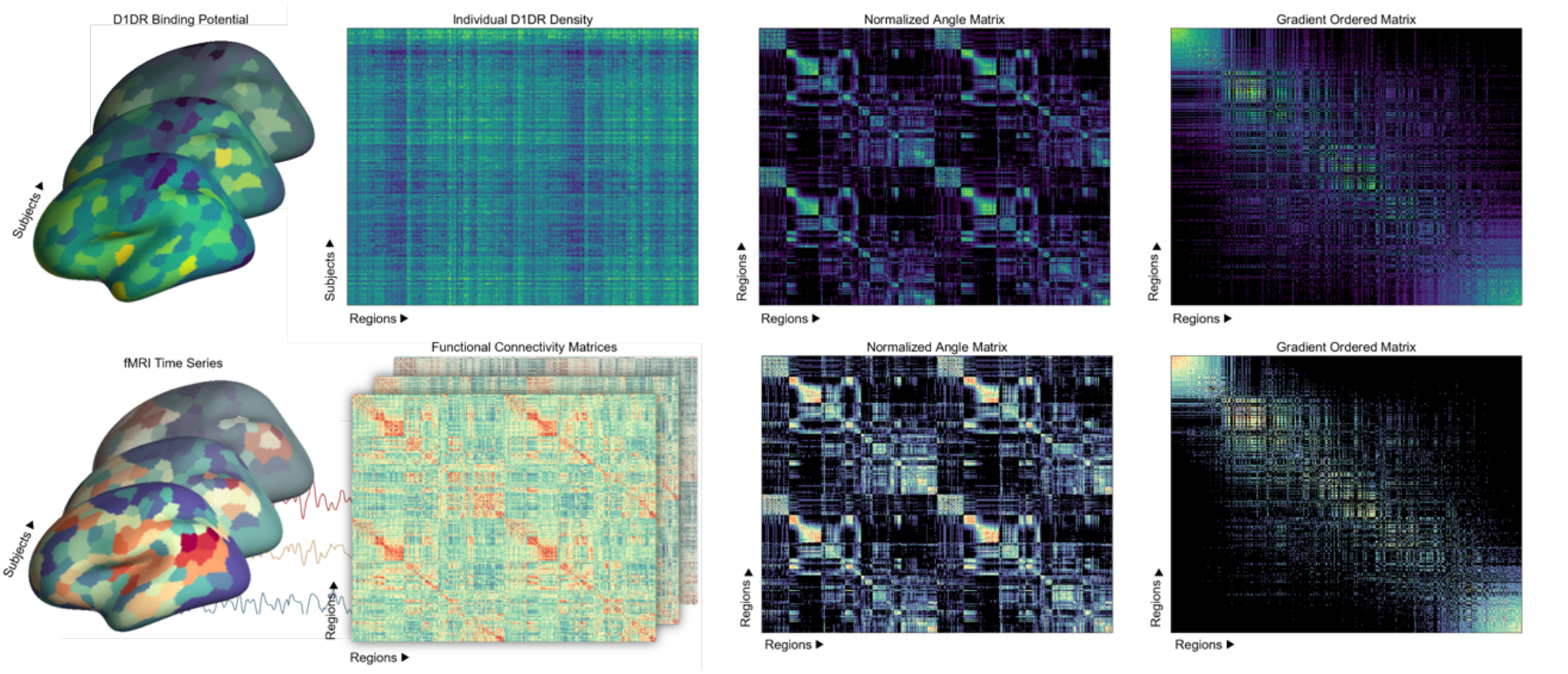
We created a group-representative cortical profile of D1DR organization and functional connectivity based on [11C]SCH-39166 PET and rsfMRI images from a total of 180 healthy participants. Top: Subject-specific binding potentials were sampled from 400 cortical parcels. Between-subject Inter-Regional Correlation Analysis was used to compute inter-regional D1DR covariance, adjusted for differences in age and sex. The group-level D1DR covariance matrix was converted to a normalized angle matrix and subjected to Laplacian eigenmapping, allowing decomposition of high dimensional affinity matrix into low dimensional representations of covariance patterns. Bottom: denoised rsfMRI time series were sampled from the same cortical parcels as described for D1DR to create subject-specific connectivity matrices. Both a group average connectivity matrix and subject-specific matrices were then subjected to the same eigenmapping approach described above. The group-level gradients were used to test multimodal correspondences while subject-specific maps were used to test for individual differences in gradient composition in relation to D1DR density.

The first four gradients explained 60.14% of the total variance, dropping to <10% for each subsequent gradient (SI Materials and Methods). Three of the first four gradients were highly similar to the convention set by previous work^19,20^, depicting differentiation in connectivity from unimodal-to-transmodal (G1), visual-to-sensory (G2) and visual-to-executive control (G3) regions (Fig. 2). To characterize cortical organization of D1DR, a group-level inter-regional covariance matrix was computed as linear correlations of D1DR binding across subjects, reflecting regional similarity in D1DR density, adjusted for individual differences in age and sex. The same Laplacian eigenmapping technique was employed to decompose cortical gradients of D1DR covariance. Procrustes alignment^32^ was used for multimodal comparisons, a method able to linearly rotate the D1DR embedding space to a group-level representation of the functional data. Given sufficiently similar embedding spaces, alignment may resolve the order and direction of eigenvectors between modalities while preserving the manifold structure.

**Fig 2.**
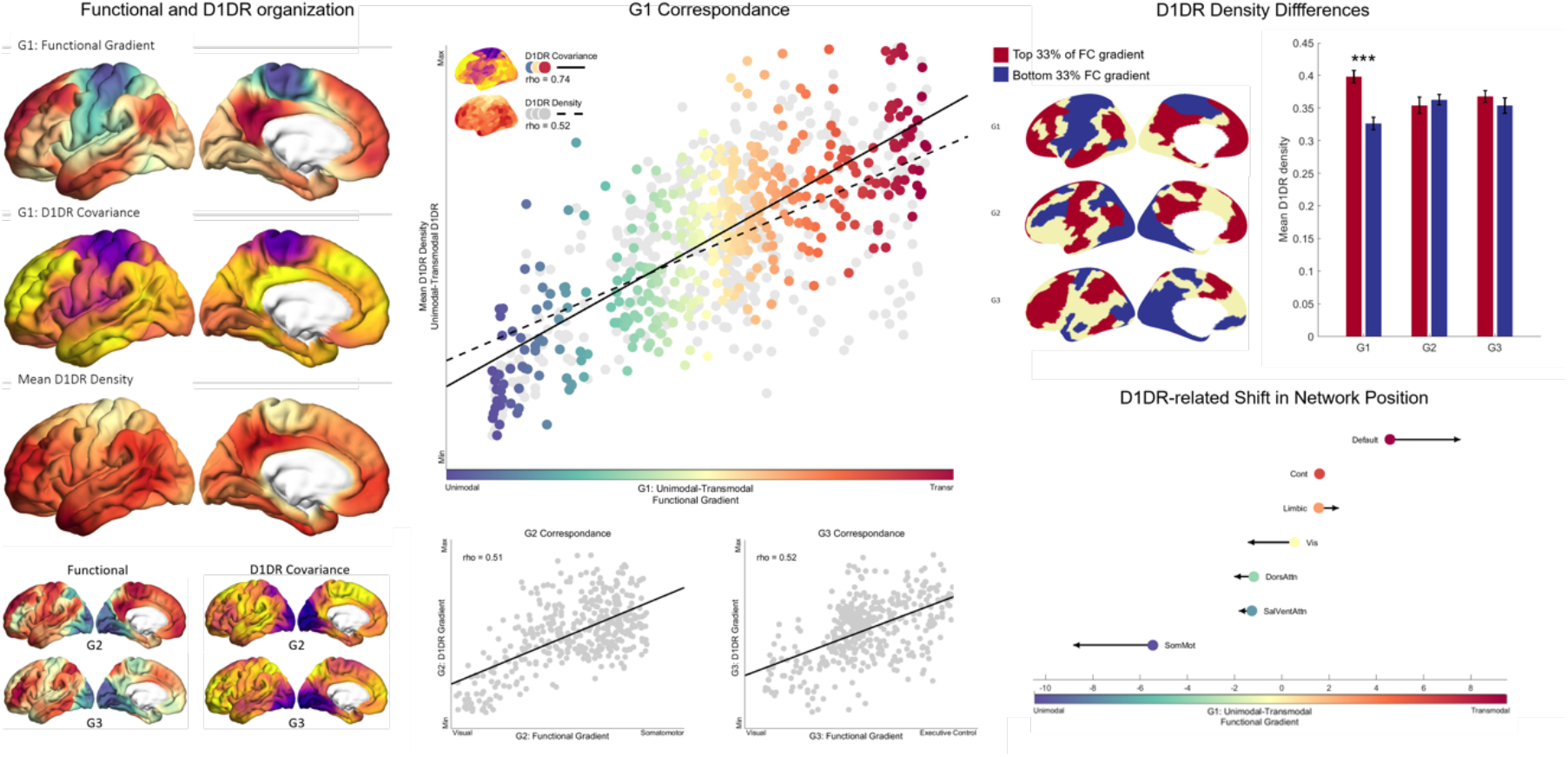
Left: Using a low-dimensional decomposition approach, we found a similar topographic profiles in both functional connectivity and D1DR covariance, depicting differentiation in connectivity from unimodal-to-transmodal (G1), visual-to-sensory (G2) and visual-to-executive control (G3) regions. Middle: We found a high spatial correspondence between D1DR covariance and the unimodal-to-transmodal axis of functional organization (G1), significantly greater compared to the cortical distribution of D1 receptors, and the respective associations of dopaminergic gradients G2 and G3. Top right: D1DR density differed significantly between unimodal and transmodal cortical regions, but not between regions corresponding to the apices of G2 and G3. Bottom right: Greater differentiation in unimodal and transmodal receptor density across individuals was associated with greater bimodality in function, reflected by greater separation between the default mode network and visual and somatomotor networks along the unimodal-to-transmodal functional axis. Colored marker placement represents the mean gradient value along the G1 axis for each network. Arrow size and direction indicate effect size of D1DR density delta between the apices of G1.

### D1DR co-expression shares organizational principles with functional architecture along a unimodal-transmodal axis

The first D1DR gradient revealed a highly similar topography to the functionally defined unimodal-to-transmodal functional gradient (G1) (rho = 0.74, *p*_spin_ < 0.001), differentiating D1DR covariance between sensory and default-mode regions. The second and third D1DR gradient expressed moderately strong correspondences to the visual-to-sensory (G2) (rho = 0.51) and visual-to-executive control (G3) (rho = 0.52) gradients, respectively (all *p*_spin_ < 0.001). Importantly, the correspondence between D1DR and FC along the unimodal-to-transmodal axis was significantly greater compared to the respective associations for G2 and G3 (F(2)=8.66, *p* = 0.0002). Further analysis revealed that the distribution of mean D1DR density expressed a similar unimodal-transmodal association (*r* = 0.52, *p*_spin_ = 0.005), although significantly weaker compared to the first D1DR gradient (*Z*_diff_ = -5.18, *p* < 0.001). Together, our results revealed dopaminergic differentiation along a unimodal-to-transmodal cortical hierarchy, both in terms of receptor covariance and density distribution, largely aligned with the principal organization of functional connectivity.

Given the role of dopamine for enhancing signal-to-noise ratio and neuronal gain^33^, we next investigated whether the shared organization varies across individuals. We hypothesized that a more pronounced hierarchal composition of dopamine receptors, i.e., greater differentiation in D1DR density between the apices of the manifold, manifest greater bimodality of function. In other words, we set out to test whether differences in D1DR density between the ends of the functional gradient is related to the position of functional networks along the functional embedding space. First, mean D1DR density only differed between the unimodal and transmodal ends of G1 (T(350) = 10.8, p < 0.001), but not between regions corresponding to either end of the G2 and G3 axes (*p*s > 0.05). We therefore limited our analysis to the unimodal-to-transmodal gradient. We proceeded by extracting a receptor density delta between regions corresponding to each subject’s unimodal and transmodal functional apices to test whether subject-specific differences in unimodal-transmodal D1DR density is associated with the relative position of functional networks on the gradient axis (Fig. 2). We found that individuals with greater difference in unimodal-transmodal D1DR density exhibited greater separation between the default-mode network in relation to somatomotor (T = 3.15, p = 0.002) and visual (T = 3.75, p < 0.001) networks. This implies that the hierarchal composition of local D1DR densities, with greater differentiation between unimodal and transmodal cortical regions, is coupled with greater segregation of corresponding functional networks.

### Topographical overlap between D1DR organization and functional architecture

Given the topographical similarity of D1DR covariation and the principal axis of macroscale functional organization, we next quantified the correspondence in D1DR covariance magnitude in relation to the functional resting-state network structure. In line with our hypothesis, mean D1DR covariance was significantly greater within functional resting-state networks compared to regions between unrelated networks (*within networks (n = 12618):* mean ± 95% CI = 0.417 ± 0.004; *between networks (n = 67182):* mean ± 95% CI = 0.348 ± 0.002; *p*_spin_ < 0.001) (Fig. 3). These results suggest a cortical distribution in D1DR covariance that is systematically aligned with functional network structures, such that functionally integrated regions exhibit greater similarity in D1DR expression than functionally segregated systems.

**Fig 3.**
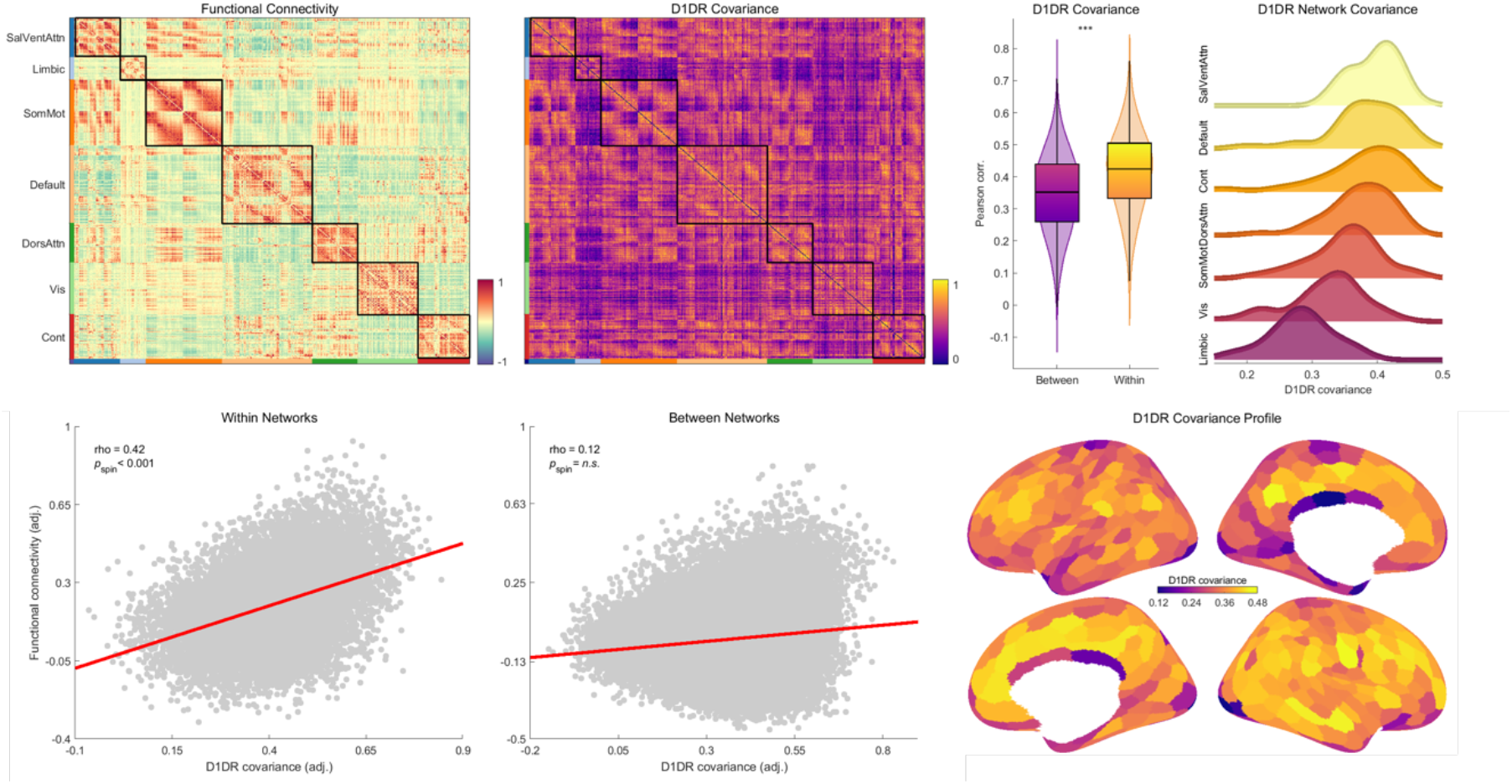
Top left: Visualization of group-level functional connectivity and D1DR covariance matrices. Inter-regional D1DR covariance was significantly greater within functionally demarcated networks compared to covariance between networks. Edgewise associations between D1DR covariance were associated with the corresponding functional connections within functional networks, but not connections between networks. Bottom right: cortical D1DR covariance profile represented by the row-wise mean of D1DR covariance. Top right: D1DR covariance profile represented by network, indicating a gradual shift in covariance from higher-order networks to lower-order sensory and limbic systems.

### D1DR organization is related to functional connectivity strength

We have demonstrated that inter-regional covariance in D1DR expression exhibits a topographically similar architecture to functional organization across unimodal and transmodal regions and functional subsystems. However, a topographical overlap does not inform whether regional similarity in D1DR expression modulate the strength of functional connections. Thus, we next tested whether the magnitude of D1DR covariance between regions is associated with the strength of corresponding functional connections. To ensure that any potential association was not biased due to spatial proximity, we controlled for linear and quadratic effects of Euclidean distance. We found that D1DR covariation was significantly related to functional connectivity within functional networks (*r* = 0.42, *p*_spin_ = 0.01), whereas no association was observed between regions related to different networks (*r* = 0.12, *p*_spin_ = 0.69) (Fig. 3). To further examine whether the difference between within and between-network associations were dependent on the magnitude of D1DR covariance, between-network edges were thresholded to closely match the mean covariance within networks (fraction of edges removed: 27.87%). Importantly, thresholding did not increase the inter-network association between D1DR covariation and FC (*r* = 0.06, Z_diff_ = 0.16, *p* = 0.98). However, while the D1DR is the most abundant DA receptor system, recent findings suggest that the degree of similarity between different neuroreceptors mediate the strength of functional couplings^8^. To test the degree of unique variance explained by D1 for intra-network connectivity, an additional test was performed while controlling for the similarity profile of 18 other receptors and transporters, in addition to Euclidean distance (see SI Methods and Materials for details). The result showed that the association between D1DR covariance and intra-network FC remained significant (*r* = 0.36, *p*_spin_ < 0.001; Z_diff_ = 4.93, *p* <0.001) even after controlling for the signature of various neurotransmitters. These results indicate that greater inter-regional similarity of the most abundant DA receptor in the cortex is associated with stronger functional connectivity within intrinsic functional systems, independent of spatial proximity, covariance magnitude, or other receptor profiles.

### D1DR and morphological organization uniquely contribute to functional architecture

The observed association between D1DR covariation and connectivity may reflect a common dependence on inter-regional differences in brain morphology (e.g., cortical thickness)^34–37^. Thus, we assessed the unique and shared covariance of functional connectivity in relation to interregional correlations in D1DR and cortical thickness, controlling for linear and quadratic effects of Euclidian distance. In line with previous reports ^34,35,38^, we observed greater covariation in cortical thickness within canonical resting-state networks compared to regions between networks (*within networks:* mean ± 95% CI = 0.210 ± 0.004 *between networks:* mean ± 95% CI = 0.174 ± 0.001 *p*_spin_<0.001). Furthermore, similar to the pattern observed for D1DRs, the magnitude of region-to-region covariation in cortical thickness was associated with the strength of functional connections within networks (*r* = 0.36, *p*_spin_ = 0.002), but not between networks (*r* = 0.06, *p*_spin_ = 0.69). Interestingly, covariation between cortical thickness and D1DR covariance expressed a moderate association both within (*r* = 0.39, *p*_spin_ < 0.001) and between (*r* = 0.35, *p*_spin_ < 0.001) networks, although differing in effect size (Z_diff_ = 3.74, *p <* 0.001). An omnibus F-test confirmed a marginal effect of both D1DR (R^2^_adj_ = 0.14, *p* < 0.001) and cortical thickness (R^2^_adj_ = 0.13, *p* < 0.001) on intra-network connectivity respectively, with only partial mediation of cortical thickness on the association between D1DR and FC (R^2^_adj_ = 0.08, Sobel test: *p* < 0.001). These results suggest a spatially homogeneous interdependence between D1DR availability and cortical thickness, in addition to unique contributions to functional intra-network connections.

## Discussion

We investigated the organization of cortical DA D1 receptor density and its relationship to functional brain architecture. We found a topographical distribution in receptor density following a unimodal-transmodal cortical hierarchy, characterized by greater density in higher-order associative regions. This is the first time a unimodal-transmodal distribution of D1DRs has been demonstrated in humans, expanding upon previous descriptions of the DA D1 system^10,39,40^. Our findings are also congruent with recent autoradiography findings, revealing a gradient in D1DR density along the macaque cortical hierarchy^18^. The discovery of a hierarchal D1DR gradient along cortical mantle can serve as a major anatomical basis by which DA modulates functional crosstalk, and in turn, relate to higher cognitive function. Further analysis revealed that the covariance structure of D1 receptors expressed greater correspondence to the principal gradient of functional connectivity compared to lower-order gradients, reflecting both functional and dopaminergic differentiation between unimodal and transmodal cortices^19,41^. Importantly, we discovered that individuals with more pronounced unimodal-transmodal receptor hierarchies exhibited greater functional differentiation between somatomotor and the default mode network. This is of particular importance, given that given that the degree of differentiation between gradient apices has been shown to decline with age and account for differences in cognitive function^42^. Taken together, our findings provide a strong support for a dopaminergic layer of brain organization, contributing to the shape of macroscale functional architecture.

Given the importance of dopaminergic modulation for inter-regional signal propagation and neural gain^9,10^, an important question is whether the chemoarchitectural organization of D1DR receptors is associated with the strength of functional couplings. To this end, we utilized a more traditional arealization approach to investigate the correspondence between inter-regional covariance in cortical D1DR density and functional connectivity. We found that the degree of D1DR covariance was significantly greater within functional brain systems compared to between systems. Critically, region-to-region covariation in receptor density was associated with the strength of functional connections within, but not between, functionally specialized subsystems. This suggests that the chemoarchitectural profile of inter-regional associations in D1DR expression is systematically aligned with functional network structure and regulate inter-regional communication. However, recent findings suggest that spatial co-expression of different receptor types is similarly coupled with the strength of functional connections^8^. While our findings are congruent with this observation, we found a unique contribution of D1DR for intra-network connectivity, independent of other receptor profiles. Notably, our measure of D1DR covariance reflects inter-regional co-expression rather than similarity between different receptor types. This is a notable difference given that different neuroreceptor profiles are likely related with distinct properties of brain function. Moreover, both DA receptor density and functional connectivity has been linked to regional gray-matter differences^34–37^. It is therefore likely that inter-regional covariation in D1DR availability is dependent on variation in cortical morphology (e.g., differences in shape, folding, and depth). Indeed, we found that region-to-region covariation in D1DR expression correlated strongly with covariation in cortical thickness, both within and between functional subsystems. Critically, both D1DR and cortical thickness uniquely contributed to the strength of functional connections within intrinsic resting-state networks. This suggests a functional dependency on distinct properties in both cortical gray-matter organization and dopaminergic receptor structure. The fact that brain regions exhibiting similar cortical morphology covary in D1DR availability points towards a structurally and chemoarchitecturally constrained axis of variability. Importantly, this axis largely overlaps with specialized functional subsystems and contributes to inter-regional signaling. This observation is in line with previous findings of structural covariance in cortical gray matter^34,35^, suggesting that shared properties in cytoarchitectonic, chemoarchitectural, and structural covariance contribute in shaping functional brain organization ^6,34,35,43–46^.

In summary, our results constitute an important step toward an understanding of dopamine D1 system organization and its role for functional brain architecture. We expand upon previous work by highlighting the importance of inter-regional relationships in D1DR expression, beyond the topographical profile of receptor availability and anatomical gray-matter properties, in shaping the functional architecture of the brain. The discovery of a dopaminergic layer of functional brain organization represents a crucial first step towards an understanding of how DA, with close ties to behavior and neuropsychiatric conditions, potentially contribute to the emergence of functional brain organization. Understanding the dopaminergic layer of functional organization will provide a potential modifiable target to ameliorating and delaying cognitive impairments using prevention and intervention strategies.

## Materials and Methods

The current study used baseline data (N = 180) from the DyNAMiC project^25^, a prospective study of healthy individuals across the adult lifespan. The study was approved by the Swedish Ethical Review Authority, and all participants gave written informed consent prior to testing. We have reported about the study’s design, imaging protocols, and procedures elsewhere^25^. Out of the 180 participants, four were excluded due to issues related with PET-imaging: one showed indications of subcutaneous injection, two were excluded due to technical problems, and one participant declined to undergo PET. The final sample for the current study included 176 participants (82 females) aged 20 – 78 years (mean = 49, SD = 17.38). Functional and D1DR gradients were created using the Laplacian eigenmapping algorithm^28^. Laplacian eigenmapping allows decomposition of the high-dimensional affinity matrices to low dimensional components while preserving local properties in the embedded space. The locality-preserving character of the Laplacian eigenmap algorithm makes it relatively insensitive to outliers and noise compared other nonlinear manifold learning techniques^26^. Gradient decomposition was performed on thresholded covariances matrices, only keeping the top 10% of row-wise edges, converted to normalized angle matrices^19^. Between-subject Inter-Regional Correlation Analysis^47^ was used to create spatial D1DR covariance matrices, controlling for linear and quadratic effects of age and differences in sex. Statistical significance for all multimodal comparisons was determined by spatial autocorrelation-preserving permutation tests (i.e., “spin-test”^38^), achieved by randomly rotating a spherical surface projection of the parcellated data 1000 times and resampled to generate a null-distribution. Additional information regarding methods is available in SI Materials and Methods.

## Acknowledgments

This work was funded by the Swedish Research Council (grant number 2016– 01936 to A.S.), Knut and Alice Wallenberg Foundation (Wallenberg Fellow grant to A.S.), Riksbankens Jubileumsfond (RJ, P20-0515 to A.S.), and StratNeuro grant at Karolinska Institutet (A.S.). We thank the staff of the DyNAMiC project, Frida Magnusson, Vania Panes Lundmark, and staff at MRI and PET labs at Umeå University Hospital, and all our participants.

