## Supplementary material for "Dopamine D1-receptor Organization Contributes to Functional Brain Architecture": SI Methods and Materials

The current study used baseline data from the DyNAMiC project [1], a prospective study of healthy individuals across the adult lifespan. The study was approved by the Swedish Ethical Review Authority, and all participants gave written informed consent prior to testing. We have reported about the study's design, imaging protocols, and procedures elsewhere [1]. Here, only methodological and material aspects of relevance for the current study is presented.

### Participants

The DyNAMiC study participants ($N$ = 180) were recruited by random selection from the population registry of Umeå, Sweden, stratified by six age cohorts between 20 – 80 years ($n$ = 30 per decade, 50% females). Participants were screened for a set of exclusion criteria, including contraindications to magnetic imaging, medical conditions or treatment potentially affecting brain function and cognition, neurological disorders, brain pathology, and cognitive impairment. All participants completed a full set of functional and structural MRI scans, and one hundred seventy seven (participants completed [$^{11}$C]SCH23390 PET scans. Four subjects were excluded due to the following: one showed indications of subcutaneous injection during PET imaging, two were excluded due to technical problems during PET, and one participant declined to undergo PET. The final sample for the current study included 176 participants (82 females) aged 20 – 78 years (mean = 49, SD = 17.38).

#### *Imaging Procedures*

Magnetic resonance (MR) imaging was conducted using a 3-tesla scanner (Discovery MR-750, General Electric), equipped with a 32-channel phased-array head coil. PET scanning was conducted using a hybrid PET/CT system (Discovery PET/CT 690, General Electric).

*PET Imaging*

Production of [$^{11}$C]SCH23390 was performed by the radiochemistry laboratory of Norrlands Universitetssjukhus, Umeå University, according to procedures described previously [1]. Injections of [$^{11}$C]SCH23390 had high molar activity and low mass (range = [205, 391] MBq, mean ± SD = 337 ± 27 MBq. Participants were positioned on the scanner bed in supine position and were individually fitted with thermoplastic masks to prevent excessive head movement. Preceding the injection, a 5-min low-dose helical CT scan (20 mA, 120 kV, 0.8 s per revolution) was obtained for PET-attenuation correction. Continuous PET-measurement in list-mode format was initiated at the time of injection and continued for 60 minutes. Offline re-binning of list-mode data was conducted to achieve a sequence of time-framed data with increasing frame length: 6 x 10; 6 x 20; 6 x 40; 9 x 60; 22 x 120 s (n = 49 frames). Time-framed, attenuation-, scatter-, and decay-corrected PET images (47 slices, 25 cm FOV, 256×256-pixel transaxial images, voxel size 0.977 × 0.977 × 3.27 mm$^3$) were reconstructed by using the manufacturer-supplied iterative VUE Point HD-SharpIR algorithm (6 iterations, 24 subsets, resolution-recovery).

Estimation of target binding potential (BP) to non-displaceable ($BP_{ND}$) binding was computed with the cerebellum as reference region [1,2]. Pre-processing included frame-to-frame head motion correction and registration to T1-weighted MRIs using Statistical Parametric Mapping software (SPM12, Wellcome Institute, London, UK). Corrected PET data were re-sliced to match the spatial dimensions and resolution of the MR-data (1 mm$^3$ isotropic voxel size, 256 x 256 x 256). Partial-volume-effect (PVE) correction was carried out using a symmetric geometric transfer matrix (SGTM; regional correction) method implemented in FreeSurfer [3], with an estimated point-spread function of 2.5 mm full-width-at-half-maximum (FWHM). Voxel-wise $BP_{ND}$ estimates were then computed using a simplified reference tissue model (SRTM[4]) and averaged in 400 cortical parcels [5]. Univariate outliers in regional D1DR estimates were identified using the box-plot method (1.5*IQR) stratified by age in decades and sex. Outliers were generally related to poor PET-model fit along the midline ($BP_{ND}$ < 0.15, 0.72% of ROIs) and replaced by age-matched means for the right and left hemisphere separately.

*MR Imaging*

High-resolution anatomical T1-weighted images were acquired by a 3D fast spoiled gradient-echo sequence. Imaging parameters were as follows: 176 sagittal slices, thickness = 1 mm, repetition time (TR) = 8.2 ms, echo-time (TE) = 3.2 ms, flip angle = 12º, and field of view (FOV) = 250 x 250 mm. Anatomical T1-weighted images were used to parcellate cortical and subcortical structures and to estimate gray-matter volume using Freesurfer 6.0 (https://surfer.nmr.mgh.harvard.edu; [6]), with manual corrections performed using the Voxel Edit mode in Freeview if necessary.

Whole-brain functional images were acquired during resting-state while subjects were instructed to keep their eyes open, let their minds wander, and remain as still as possible. Functional images were sampled using a T2*-weighted single-shot echo-planar imaging (EPI) sequence, with a total of 350 volumes collected over 12 minutes. The functional sequence was sampled with 37 transaxial slices; slice thickness = 3.4 mm, 0.5 mm spacing; TR = 2000 ms, TE = 30 ms, flip angle = 80º, and FOV = 250 x 250 mm.

The functional images were further processed to reduce artefactual influence of non-neuronal sources. All images were first corrected for slice-timing differences, motion, and signal distortions. The time-series were subsequently demeaned and detrended, followed by simultaneous nuisance regression and temporal high-pass filtering (0.008 Hz) as to not re-introduce nuisance signals [7]. Nuisance regression of physiological variables included average white matter and cerebrospinal fluid time series, six motion parameters, in addition to derivatives, squared, and squared derivatives of each variable [8,9]. To further control for motion, a set of binary spike regressors were included for volumes exceeding a relative root-mean-squared displacement of 0.5 mm or FD of 0.2 [10]. Nuisance-regressed images were normalized to a sample-specific group template (DARTEL[11]) and spatially smoothed using a 6-mm FWHM Gaussian kernel and affine-transformed to stereotactic MNI-space.

Functional connectivity graphs were created by sampling average BOLD timeseries from the same 400 cortical parcels as used for the PET D1DR estimates [5], and each region was labeled by network affiliation[12]. Subject-wise adjacency matrices were computed using Pearson's correlations followed by Fisher's r-to-z transformation and averaged across the sample for population-level analysis. The group-average graph was then transformed by an inverted hyperbolic tangent to re-normalize the range of correlation coefficients (-1 to 1) before gradient decomposition.

### Statistical Analyses

#### *Inter-Regional Correlation Analysis*

Between-subject Inter-Regional Correlation Analysis (IRCA[13]) was used to investigate the functional organization of D1-like DA receptors (D1DR). PET IRCA utilizes regional covariance in ligand uptake across subjects to characterize topological properties of molecular brain markers, with the assumption that brain regions with significantly correlated binding potentials reflect biologically meaningful connections between spatially distributed regions [13,14]. IRCA produces robust network metrics with high test-retest reliability [14], even with relatively modest sample sizes [13,15]. This approach has been used to investigate metabolic connectivity[16–23] and to characterize organization of serotonin receptors [24], transporters [25], dopamine synthesis capacity [26,27], D1DRs [28,29], and D2DRs [30–32].

The method of cross-correlating ligand binding from different regions across subjects is sensitive to inter-individual differences [14]. To reduce ancillary covariance related to age or sex-differences, least squares regression was used to estimate linear and quadratic age-effects of D1DR binding across subjects, including linear effects of sex, for each ROI. Standardized residuals were then used to compute pairwise inter-regional D1DR BP correlations followed by Fisher's r-to-z transformation, effectively yielding a ROI × ROI population-level adjacency matrix. This method is similar to using pair-wise partial correlations of D1DR BP while controlling for age and sex. To evaluate the putative effect of age-related differences on the inter-regional correlations, a separate matrix was computed with standardized D1DR BP values without age regression. The age-preserved and age-regressed adjacency matrices correlated highly (Spearman's rho = 0.786), and the age-attributable dissimilarity between the two matrices accounted for 9.2% of the total variance Consistent with previous reports [29], this indicates that age-related differences have a relatively small effect on inter-regional D1DR covariance.

*Laplacian eigenmap construction*

To compute gradients of D1DR and functional connectivity organization, we employed Laplacian eigenmapping [33], a nonlinear manifold learning technique able to identify principal gradients components. In line with previous studies [34,35], the adjacency matrices (D1DR covariance and functional connectivity) were thresholded row-wise, only keeping the top 10% of positive edges. Thresholded and negative edges were set to zeros. The matrices were then converted to normalized angle matrices. Laplacian eigenmapping was performed using the algorithm provided by Brain Space toolbox [36] for MATLAB with default settings for the main analysis. In brief, the algorithm estimates a low-dimensional embedding of the high-dimensional affinity matrices while preserving local properties in the embedded space. The locality-preserving character of the Laplacian eigenmap algorithm makes it relatively insensitive to outliers and noise compared other nonlinear manifold learning techniques [33]. The approach results in a number of eigenvectors, referred to as "gradients". To ensure the spatial correspondence of gradients between the fMRI and PET data, D1DR gradients underwent Procrustes linear alignment [37], a method that resolves the order and sign of the eigenvectors, but preserves the structure of the gradients. To ensure that the size of the rotated embedding space did not artificially inflate correspondence, control analyses were performed to confirm that the fewest number of gradients, cumulatively accounting for >50% of the total variance (n=4), yielded similar results (G1, rho = 0.65; G2, rho = 0.51; G3, rho = 0.22; following the naming convention set in the main text). Although only the third gradient (Z = 5.16, p < 0.001) expressed statistically weaker bimodal correspondence in the reduced manifold space.

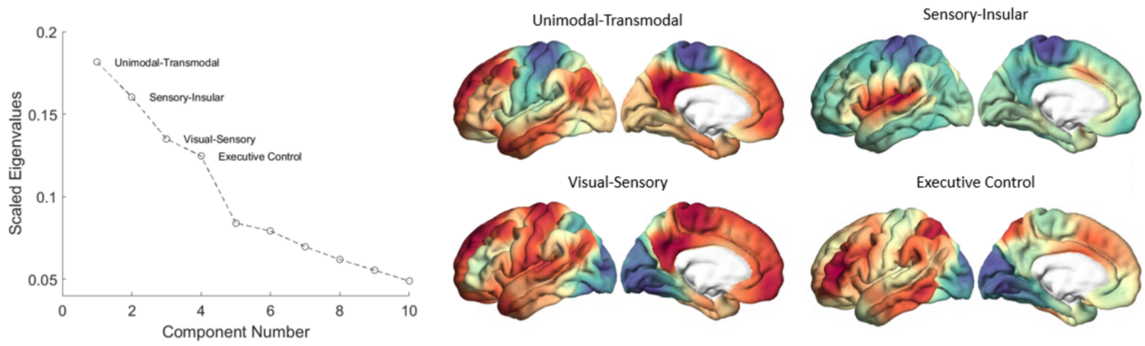


**SI. Fig. 1.** Scree plot depicting degree of variance explained by the first 10 gradients. The top four gradients explained 60.14% of the total variance. Three out of the four top gradients were highly similar to the convention set by previous work (Unimodal-Transmodal; Visual-Sensory, Visual-Executive Control) and used for the main analyses.

*Association between individual differences in D1DR density and network position*

We defined the apices of each gradient by creating binary maps of the upper and lower 33% of gradient values for each participant. Mean D1DR density was extracted from regions in the subject-specific binary maps of gradient apices. A D1DR density delta was computed as the absolute difference between the upper and lower gradient percentiles. Placement of resting-state networks[12] along the functional manifold was carried out as described in ref.[38]. In brief, the center of mass of each network was first computed as the median gradient value of network parcels. Distances between each network's center of mass was then computed as the pairwise difference between networks. For each network pair, linear regression models were fitted with subjects' network distances as dependent variable, and D1DR density delta as independent variable. Each model included variables of subjects' age, sex, and FD as nuisance variables of no interest. Statistical significance was determined using FDR-corrected p-values.

*D1DR covariance within and between functional systems*

To investigate whether inter-regional D1DR covariance corresponds to the modular system-level topography of the functional connectome, we expected a difference in D1DR covariance within and between canonical resting-state networks. Moreover, we hypothesized that the topological profile of D1DRs covaries with inter-regional differences in FC and cortical thickness between functionally coupled brain regions. To test these possibilities, all cortical parcels were labeled according to their network affiliation as defined in a seven-network atlas[12]. To examine the difference in intra- and inter-modular D1DR covariance and cortical thickness, we used spatial autocorrelation-preserving permutation tests (i.e., "spin-test" [39]). This was achieved by first creating surface-based representations of all subjects' D1DR biding potential maps on Freesurfer's fsaverage surface, in addition to using surface-based thickness estimates from Freesurfer. Next, a spherical projection of the 400 parcel Schaefer atlas was randomly rotated 1000 times. For each rotation, subjects' surface based D1DR BP values were extracted for the rotated parcels and new adjacency matrices were computed as described previously. A null distribution of intra- and inter-modular edges were subsequently computed based on the surface-rotated matrices.

*Edgewise associations between D1DR, FC, cortical thickness, and other receptor profiles*

Associations between inter-regional D1DR correlations, FC, and cortical thickness were assessed by partial correlations (Spearman's rho) of coaxial intra- and inter-network edges, respectively, controlling for linear and quadratic effects of Euclidean distance between ROIs. To this end, IRCA was used to compute a population-level cortical thickness adjacency matrix. First, ROI-wise surface-based cortical thickness estimates were adjusted for sex and linear and quadratic age effects using the method described for D1DR correlations. We further performed additional analyses to test the specificity of D1DR covariance in relation to functional connectivity by controlling for the 18 other receptor profiles available in the NeuroMaps toolbox [40], excluding D1DR. Parcellated mean receptor maps (400 cortical parcels [5]) were weighted and spatially correlated following the method outlined in ref.[41], yielding a receptor-similarity correlation matrix. Intra-network receptor-similarity edges were subsequently used as a covariate in addition to linear and quadratic effects of Euclidean distance. Statistical significance of partial correlations was assessed by spin-testing, randomly rotating a spherical projection of the parcellation maps 1000 times and two-tailed statistical significance was determined at a 95% confidence level.
